# High-Coverage Four-Dimensional Data-Independent Acquisition Proteomics and Phosphoproteomics Enabled by Deep Learning-Driven Multi-Dimensional Prediction

**DOI:** 10.1101/2022.06.12.495786

**Authors:** Moran Chen, Pujia Zhu, Pengfei Wu, Yanhong Hao, Zhourui Zhang, Jian Sun, Wenjing Nie, Suming Chen

**Affiliations:** The Institute for Advanced Studies, Wuhan University, Wuhan, Hubei 430072, China

## Abstract

Four-dimensional (4D) data-independent acquisition (DIA)-based proteomics is an emerging technology that has been proven to have high precursor ion sampling efficiency and higher precursor identification specificity. However, the current 4D DIA proteomics is still dependent on the building of project-specific experimental library which is time-consuming and limits the coverage for identification/quantification. Herein, a workflow of 4D DIA proteomics by using the predicted multi-dimensional in silico library was established. A deep learning model Deep4D that could high-accurately predict the CCS and RT of both the unmodified and phosphorylated peptides was developed. By using an integrated 4D in silico library containing millions of peptides, we have identified 25% more protein than using experimental libraries in the DIA proteomics analysis of HeLa cells. We further demonstrate that the introduction of in silico prediction library can greatly complement the experimental library of directly obtained phosphorylated peptides, resulting in a greater increase in the identification of phosphorylated peptides and phosphorylated proteins.

## Introduction

Mass spectrometry (MS)-based proteomics has become an indispensable tool in the studies of life sciences and related fields. Although it has enjoyed tremendous success, MS-based proteomics still faces significant technical challenges. The conventional proteomics based on data-dependent acquisition (DDA) is often limited by proteome coverage and reproducibility due to the compromised MS sequencing speed and semistochastic selection of the precursor ions. The emerging data-independent acquisition (DIA)-based proteomics is a promising approach that could overcome these limitations by unbiased MS/MS analysis of all peptide ions within defined isolation windows.^1, 2^ However, data mining for DIA is extremely difficult since the fragments of various precursor ions may present on a single MS/MS spectrum, and most of the current DIA proteomics methods need a high-quality spectral library containing retention time (RT) and fragment ions intensities of each peptide to be constructed prior to data processing.^3^ Typically, a sample-specific DDA library was built through DDA analysis of extensively pre-fractionated or repeatedly injected samples.^4^ This approach is not only time-consuming, but also limits the identification range of peptides and reduces the coverage of protein identification. To solve this problem, several methods based on deep learning have been developed to perform DIA proteomics based on predicted in silico libraries containing RT and fragment ions intensities,^5^ such as DeepDIA,^4^ Prosit,^6^ DeepMass:Prism,^7^ and Deepphospho.^8^ Despite these progresses, the prediction accuracy, identification reliability, and proteome coverage in DIA proteomics still need to be improved.

Four-dimensional (4D) proteomics is an emerging technology which adds a fourth dimension, ion mobility, on top of the conventional proteomics that based on the three dimensions of RT, mass-to-charge ratio (m/z), and ion intensity to identify and quantify peptides. The addition of ion mobility separation to the chromatographic and mass separation increases sensitivity and reduces spectral complexity.^9, 10^ More importantly, the inherent collisional cross section (CCS) value of peptides derived from the measured ion mobility could enhance the reliability of their identification^11^. 4D DIA-based proteomics has been proven to have high precursor ion sampling efficiency and higher precursor identification specificity.^12^ Nevertheless, the current 4D DIA proteomics is still dependent on the building of project-specific experimental library. Recently, the prediction of the CCS of unmodified peptide was conducted based on long short-term memory (LSTM),^13^ which took the first step of 4D DIA proteomics based on predicted library. However, the building of the integrated in silico library containing more accurate CCS and RT required for 4D DIA proteomics and phosphoproteomics remains a great challenge.

Here, a workflow of 4D DIA proteomics by using the predicted multi-dimensional in silico library was established (Fig. 1a). A deep learning model Deep4D that could high-accurately predict the CCS and RT of both the unmodified and phosphorylated peptides was developed, which is the model based on self-attention^14^ that completely avoid the use of recurrent neural network (RNN) or LSTM (Fig. 1b). Deep4D exhibited higher accuracy in the prediction of CCS and RT of peptides than the current models based on deep learning. By using Deep4D and MS/MS prediction tool, an integrated in silico library containing CCS, RT, and fragment ion intensities of millions of peptides for 4D DIA proteomics was established based on the SwissProt H. sapiens database, which enables the deeper peptide and proteome coverage for human samples compared to using sample-specific experimental library. We further demonstrate that the introduction of in silico prediction library can greatly complement the experimental library of directly obtained phosphorylated peptides, resulting in a greater increase in the identification of phosphorylated peptides and phosphorylated proteins.

**Fig. 1.**
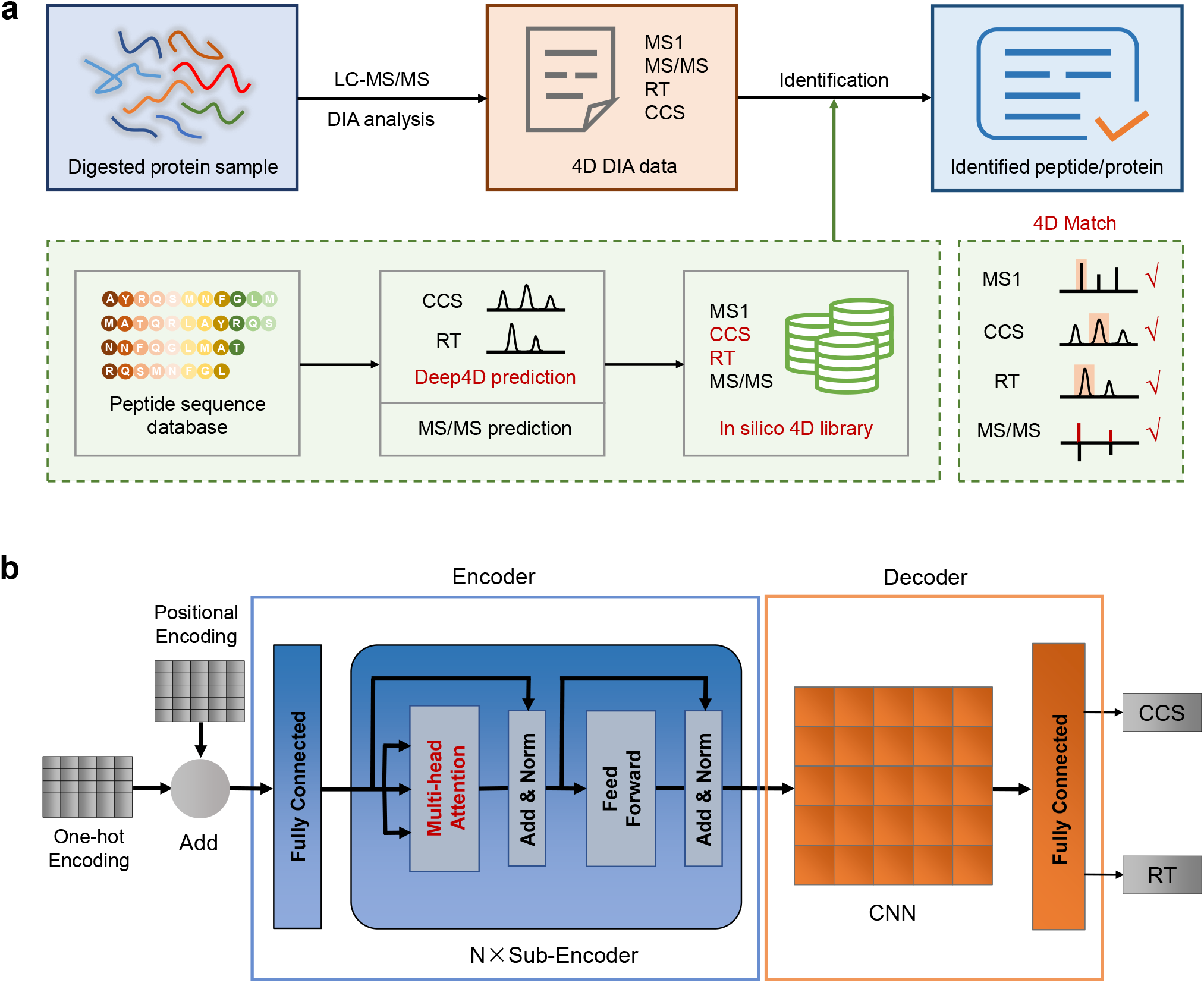
Schematic diagram of 4D proteomics and deep learning model. **a** Workflow of data-independent acquisition (DIA) proteomics based on in silico 4D peptide library. **b** Structure and mechanism of Deep4D model for peptide CCS and RT prediction.

## Results

### Development of Deep4D model and its validation in CCS value prediction

To realize the 4D DIA proteomics, the peptide-centric approach was adopted due to its power in resolving highly complicated DIA data (Fig. 1a). To accurately predict the CCS values and retention time (RT) of peptides for 4D in silico library, a Deep4D model based on the self-attention mechanism that adopts the encoder-decoder structure^15^ was developed (Fig 1b). Different from the sequence to sequence method in natural language processing^16^, the output of Deep4D changes from sequence to one constant. To gain a deep representation from peptide sequence without LSTM, multi-head self-attention was used as the core of the encoder. Compared with LSTM which aligns the positions to steps in computation time, self-attention could focus more on the global information of the sequence and be better parallelized. Convolutional neural network (CNN) and multilayer perceptron (MLP) were chosen to serve as the decoder to convert the representation of peptide to CCS or RT.

To evaluate the performance of Deep4D in predicting peptide CCS, we firstly compared Deep4D with the state-of-the-art bi-LSTM model proposed by Meier et al.^13^ Both models were trained and tested on the large-scale peptide CCS dataset (Pep_CCS, *n* = 718,884) built in that study (Fig. 2a).^13^ Instead of mixing all charged peptides, we used transfer learning to train the model with peptides of different charge states separately (*n* = 533,636). Deep4D was firstly trained on the training set of doubly charged peptide training set, then the models were fine-tuned with triply charged peptides and quadruply peptides, respectively. To avoid the over-fitting of the model, an independent additional dataset from the synthetic ProteomeTools peptides^13, 17^ was used as an external test set (*n* = 185,248). In this test set, our model shows an overall high accuracy with only a 1.25% median relative error (MRE, compares with the experimental value) (Fig. 2b and 2c), which is 11% lower than that obtained from bi-LSTM (1.4%). For the subset of peptides with different charge states, the MRE of CCS values predicted by Deep4D were also lower than those from bi-LSTM (Fig. 2d). The accuracy for predicting the doubly charged peptides could even reach a 1.03%

**Fig. 2.**
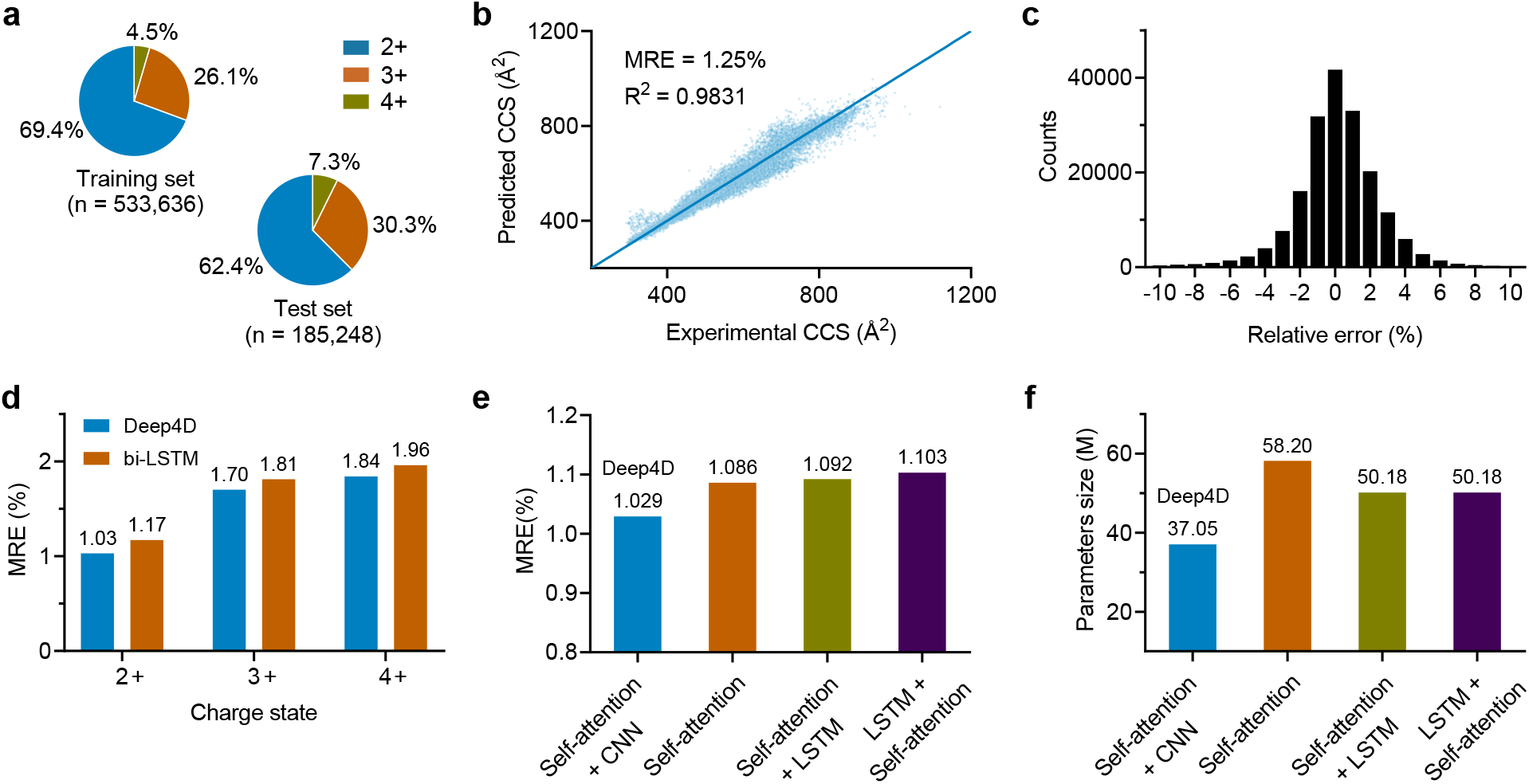
Performance of Deep4D model for the CCS prediction of peptides. **a** Information of the large-scale peptide CCS dataset (Pep_CCS). **b** Correlation of predicted versus experimental CCS values (*n* = 185,248). Correlation coefficient of linear regression (R^2^) and median relative error (MRE) are indicated. **c** Distribution of the relative errors of predicted CCS values. **d** Comparison of the median relative error (MRE) in peptide CCS prediction between the Deep4D and bi-LSTM models. **e** Comparison of MRE in peptide CCS prediction among different models. **f** Comparison of parameter sizes among different models.

The superiority of our model is not only reflected in the addition of self-attention module, but also in the overall structural design. We compared Deep4D with three other models (Supplementary Fig. 1) which contained self-attention module and used MLP in the last part of the model (Fig. 1e and 1f). Four models were tested on the doubly charged peptides derived from PepCCS dataset (Supplementary Fig. 2). The MRE of predicted CCS values obtained from these models are all lower than those from the bi-LSTM model (Fig. 1d), which indicate the benefits of using self-attention to represent peptide. Interestingly, the combination of bi-LSTM with self-attention performed worse than the model with only self-attention. Among these models, Deep4D shows the best performance with the least parameters (Fig. 1f). These results show that Deep4D is a very promising deep learning model, showing better results than previous models in CCS value prediction of peptides.

### Optimization of the Deep4D model for predicting the CCS values of phosphopeptides

Besides the unmodified peptides, the prediction of CCS values for phosphopeptides is also essential, especially for the 4D DIA-based phosphoproteomics. However, phosphorylated peptides have a more complex structure, and the modification of phosphate at different sites increases the difficulty of prediction. Thus, the high-quality phosphorylated peptide CCS dataset and the prediction tools are still lacking. To this end, we try to establish a large-scale phosphopeptide CCS dataset across species to optimize Deep4D, and empower its ability to accurately predict CCS of phosphopeptides.

To build this dataset, we first extracted and enriched phosphopeptides from different species, including cell lines of HeLa and HepG2, tissues of mice brain, kidney, and liver, and yeast. Two different proteolytic enzymes (trypsin and chymotrypsin) were used to digest proteins to generated more peptides with diverse cleavage sites. After the subsequent LC-MS/MS analysis in trapped ion mobility spectrometry (TIMS) mass spectrometer and peptide identification, we constructed a 4D phosphopeptide library containing 54,125 phosphopeptides with highly-confident phosphorylation site information (Fig. 3a and 3b, Supplementary Table 1-3). The C-terminus of phosphopeptides contains a wide range of amino acids, which makes the peptide structure more comprehensive. The results show that the doubly (82%) and triply (18%) charged phosphopeptides are predominant, and most of the peptides contain one or two phosphate groups (Fig. 3b). Across the 20,536 phosphopeptides measured at least in duplicate, the median coefficient of variation (CV) was only 0.35%, which highlights the excellent reproducibility of TIMS CCS measurements (Fig. 3c).

**Fig. 3.**
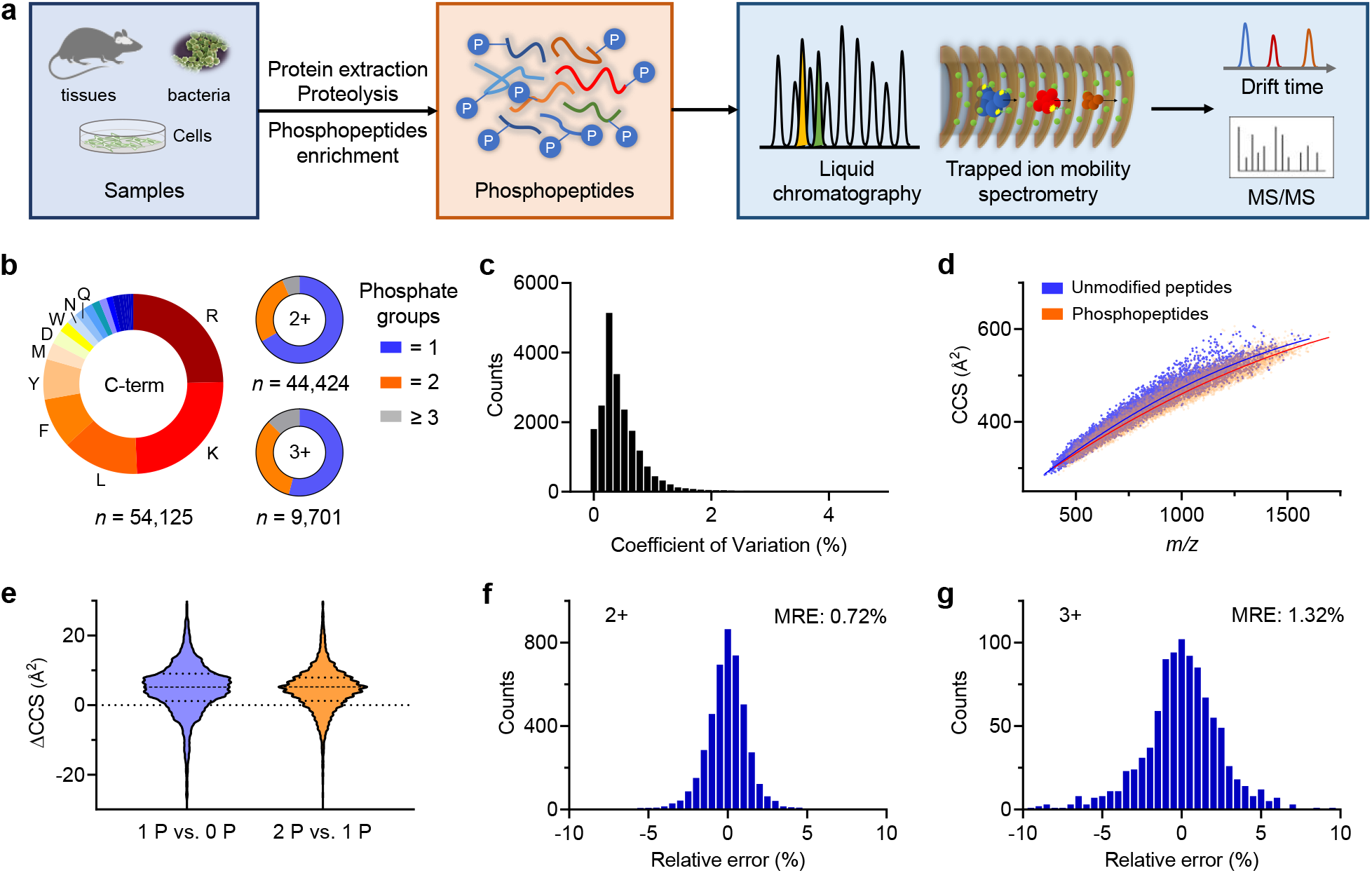
Development of Deep4D model for the prediction of phosphopeptide CCS. **a** Schematic diagram of the construction of large-scale phosphopeptides library. **b** Information of the phosphopeptides in the dataset. **c** CVs of repeatedly measured phosphopeptide CCS values in the full dataset (*n* = 54,125). **d** Distribution of the unmodified peptide and phosphopeptides data points in the CCS vs. *m/z* space. Trend lines (blue and red) are fitted by second order polynomial. **e** The distribution of CCS difference of each peptide pair. ΔCCS of each pair was calculated by subtracting CCS of unmodified peptide (0P) from mono-phosphosite (1P) peptide (blue) and subtracting CCS of mono-phosphosite (1P) peptide from dual phosphosite peptide (2P) (orange). **f, g** Relative errors in the prediction of CCS values for phosphopeptides with (**f**) 2+ and (**g**) 3+ charge states using Deep4D model.

Having established the relatively large-scale dataset for phosphopeptides with CCS values, we explore whether a dataset of this scale could shed light on the influence of phosphorylation on the change of peptide CCS. Plotting the mass-to-charge (*m/z*) vs. CCS distribution of doubly charged unmodified peptides and phosphopeptides separates them over the *m/z* range 350-1700 Da and 285-600 Å in collisional cross section (Fig. 3d). The trend line of phosphopeptides appear below that of the unmodified peptides, which indicates the phosphopeptides have more compact conformations than the peptides with the same molecular weight. The results imply that phosphate moiety may form intramolecular interactions with the neighboring residues and peptide backbone, resulting in relatively compact structures. The consistent result was also observed in a previous study obtained in a small-scale dataset.^18^ To further investigate the effect of modification of phosphate groups on the CCS of peptides, we selected 2948 pairs of unmodified and monophosphorylated peptides with the same sequence from the dataset and counted the difference between the CCS of phosphorylated and unmodified peptides (Fig. 3e). The results showed that 78% (*n* = 2291) of the differences were greater than zero and only 22% (*n* = 657) of the differences were less than zero. Similarly, we compared the difference in CCS values between 2581 pairs of bisphosphates modified and monophosphate modified peptides, where 79% (*n* = 2035) of the peptides had CCS differences greater than zero and 21% (*n* = 546) of the peptides had CCS differences less than zero. The above results indicate that if additional modified phosphate groups are added to the peptide, it will make the CCS value of most of the peptides larger.

Based on the established 4D dataset of phosphopeptides, the Deep4D model was further fine-tuned to fit the prediction of the CCS values of phosphopeptides. The training and prediction of the doubly charged (*n* = 44,424) and triply charged (*n* = 9701) phosphopeptides were conducted separately. The subset of each charge state is randomly divided into a training set and a test set according to the ratio of 90% and 10%. In the test set of doubly charged phosphopeptides (*n* = 4446), our model reached a high accuracy with an MRE of 0.72% (Fig. 2f). The test set of triply charged phosphopeptides (*n* = 971) gave an MRE of 1.32% (Fig. 2g). These results show that Deep4D also performs very well in predicting the CCS values of phosphopeptides.

### Accurate prediction of retention time for peptides with Deep4D

Liquid chromatography (LC) is a widely used separation method in MS-based proteomics, and the retention time (RT) is also a key dimension in DIA proteomics. Given that the Deep4D model has shown superiority in predicting the CCS values of peptides, we reason that it will also perform well in predicting the RT of peptides. To verify the speculation, Deep4D was compared with other state-of-the-art models for RT prediction of both unmodified and phosphorylated peptides.

Firstly, we compared Deep4D with DeepLC on RT prediction of unmodified peptides. DeepLC is a convolutional deep learning model with peptide encoding based on atomic composition and performs similarly to other state-of-the-art approaches for unmodified peptides.^19^ To avoid possible unfairness when retraining this model with new datasets, we use the 15 unmodified peptide datasets mentioned in their work for comparison. These datasets were represented with experimental RT or indexed RT (iRT).^20^ The Deep4D model was fine-tuned with each specific dataset before the prediction. In terms of mean absolute error (MAE) (Fig. 4a and 4b) and Δt95% (the minimal time window containing the deviations between observed and predicted RTs for 95% of the peptides, Supplementary Fig. 3 and 4), Deep4D outperformed DeepLC in 14 datasets which contain three types of LC data [reversed-phase LC (RPLC), hydrophilic interaction LC (HILIC), and strong cation exchange (SCX) chromatography], except for HeLa HF dataset (0.35% vs. 0.31%, MAE). Note that HeLa HF is built through a substantially shorter LC gradient, which could cause a negative influence on resolution and peak capacity, and make apex peptide RT less predictable.^21^ Fig. 4c and 4d show the correlation between the experimental observations and the Deep4D predicted values for the two selected datasets. We observe very high prediction accuracy, with Pearson correlation coefficients larger than 0.99 for both datasets. These results suggest that Deep4D is overall better at predicting the RT of unmodified peptides.

**Fig. 4.**
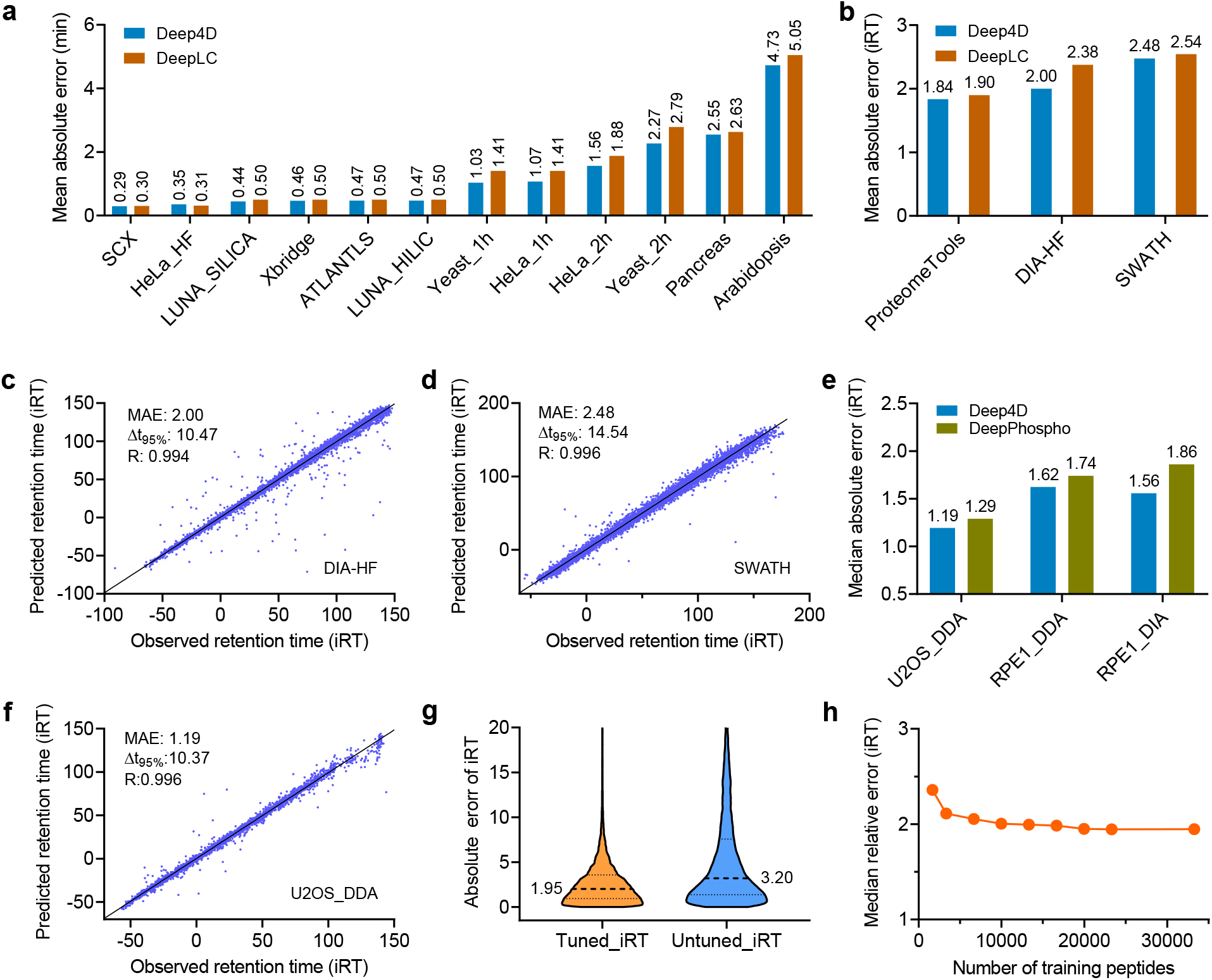
Performance of Deep4D on RT prediction. **a, b** Comparison of the prediction accuracy for (**a**) RT and (**b**) iRT of peptides with Deep4D and DeepLC models in term of the mean absolute error on different datasets. **c**, **d** Correlation of predicted versus experimental iRT values of peptides with Deep4D on (**c**) DIA-HF and (**d**) SWATH datasets. **e** Comparison of the prediction accuracy for iRT of phosphopeptides with Deep4D and DeepPhospho models in term of the median absolute error on different datasets. **f** Correlation of predicted versus experimental iRT values of phosphopeptides with Deep4D on U2OS_DDA dataset. **g** Distribution of absolute errors of peptide iRT predicted by fine-tuned and untuned Deep4D model. **h** Relationship between the number of training peptides for Deep4D and its prediction accuracy.

We further evaluate the performance of Deep4D on RT prediction of phosphopeptides by comparing it with DeepPhospho, which is the only model that uses self-attention besides Deep4D and shows great performance in predicting the RT of phosphopeptides.^8^ Likewise, Deep4D was compared with DeepPhospho on three phosphopeptide RT datasets used in their study. The results show that Deep4D outperformed DeepPhospho in all three datasets in terms of MAE (Fig. 4e). The high accuracy with a Pearson correlation coefficient of 0.996 was observed between the predicted and observed iRT in U2OS_DDA dataset (Fig. 4f).

In the prediction of the peptide RT, a model that can be fine-tuned according to instrument status and different instruments could yield a higher degree of accuracy, because RT and even iRT may be influenced by types of instruments and changes in chromatographic settings.^4, 22^ To evaluate the performance improvement of the Deep4D model after fine-tuning, we first pre-trained Deep4D with SWATH library and fine-tuned the model using a dataset of HeLa cells (HeLa1, 1-h gradient), and then tested its prediction performance on another independent dataset derived from mouse brain tissue (Brain1, 2-h gradient). These two datasets were acquired by the same LC-MS instruments and conditions. When using all the data in Hela1 (*n* = 33,263) to fine-tune Deep4D, the absolute median error of iRT prediction after fine-tuning (1.95) was 39% lower than that of untuned model (3.20) (Fig. 4g). This result shows that the tunable model has a great advantage in improving the prediction accuracy. Interestingly, we found that a small training set is enough for fine-tuning Deep4D to achieve a good performance. The use of 3,270 peptides could reduce the median error of iRT prediction to 2.1%, which is only 7.7% larger than that obtained with 33,263 peptides (Fig. 4h). In CCS prediction, we also found a large gap between the performance of fine-tuned and untuned model and Deep4D could also be easily fine-tuned with a small training set (Supplementary Fig. 5). The above results demonstrated that Deep4D is an easily fine-tunable model for accurate prediction of RT and CCS of both the unmodified and phosphorylated peptides.

### High-coverage 4D DIA proteomics enabled by constructing large-scale in silico library with Deep4D

To realize the 4D DIA proteomics that does not rely on the experimental library, the higher-coverage predicted 4D in silico library containing the information of CCS, RT and MS/MS is necessary (Fig. 1a). Deep4D has shown the ability to accurately predict CCS and RT of peptides. We next want to combine the accurate MS/MS prediction tool DeepPhospho^8^ to generate the 4D predicted library and test how this approach will perform in DIA proteomics (Fig. 5a).

**Fig. 5.**
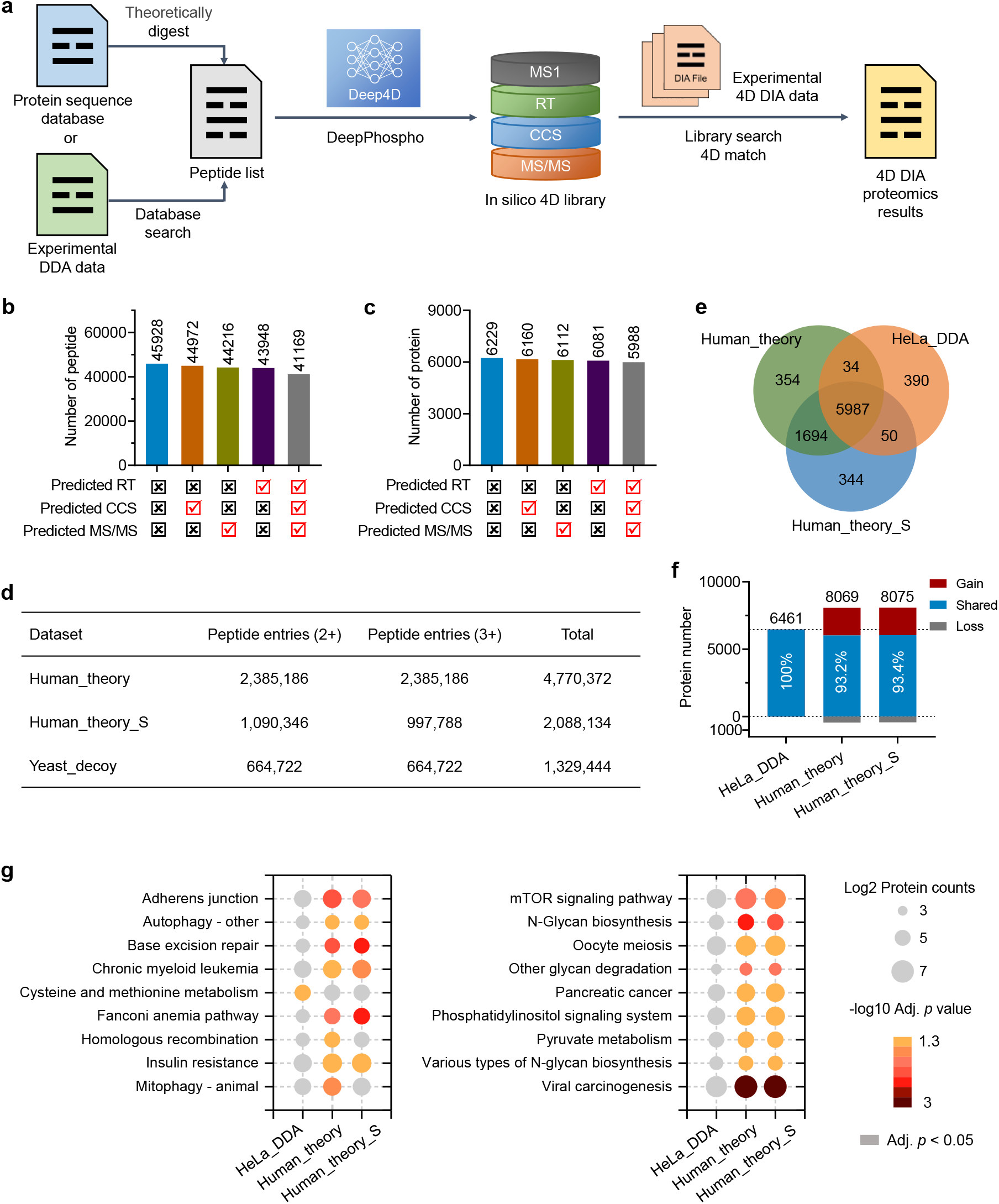
DIA-based proteomics driven by in silico 4D libraries. **a** Workflow for the construction of in silico 4D libraries and the application in DIA proteomics. **b, c** Comparison of the identified numbers of (**b**) peptides and (**c**) proteins in DIA proteomics analysis of HeLa cells with different 4D libraries. **d** The peptide information of different 4D libraries. **e** Venn diagram of the identified proteins in DIA proteomics analysis of HeLa cells with different 4D libraries. **f** Comparison of the identified proteins between the using of full predicted libraries and experimental library. **g** The different enriched KEGG pathways based on proteome profiling results yielded between the experimental HeLa_DDA library and predicted libraries. Fisher’s exact test (two-sided) was performed and adjusted by the Benjamini–Hochberg procedure. Significantly enriched pathways (adjusted *p* < 0.05) are annotated in a color gradient, and enrichment terms with adjusted *p* > 0.05 are shown in light gray.

The validation was first performed for identifying the proteins in HeLa cells by using an in silico 4D library derived from the experimentally acquired peptide library. The initial peptide sequences were acquired by analyzing the peptides from a separate HeLa cells sample with timsPro mass spectrometer using DDA method, and then the CCS, RT values and MS/MS spectra were predicted by Deep4D and DeepPhospho, which were fine-tuned with peptides information from RAW-264.7 cells (Supplementary Fig. 6). To comprehensively evaluate the performance of the predicted library, we created 4D peptide libraries by replacing one or all the information of CCS, RT and MS/MS in the experimental library with predicted values. As shown in Fig. 5b, the numbers of the identified peptides decrease a little bit if the predicted values were used in the libraries, but only a 10% reduction was observed (from 45,928 to 41,169) even with all predicted values. For the identification of proteins, the total number only decreases by 5 % when using the all-predicted library (from 6229 to 5988, Fig. 5c). A slight reduction in the number of peptide and protein identifications is reasonable when using prediction libraries, which are derived from the experimental library. These results indicate that the performance of predicted 4D library is almost comparable to the experimental library.

Building a prediction library from an experimental library for each sample is not the most efficient route; sample-specific libraries are not only time-consuming, but also limit the size of the prediction library and thus the number of identified proteins. To overcome this limitation, we used all reviewed proteins (20,387 proteins) from SwissProt H. sapiens database to build a more widely applicable in silico peptide library. The peptide list was generated by tryptic specific digestion with missed cleavage set to 2 and every peptide was given charge states 2+ and 3+. The in silico 4D library (Human_theory) containing 4,770,372 peptide entries is constructed by Deep4D and DeepPhospho with the same parameters as before and is expected to be applied to a variety of human samples without the need for experimentally library building for each sample (Fig. 5d).

Then, this large-scale Human_theory 4D library was applied to the DIA proteomic analysis of HeLa cell sample. The identified proteins were compared with that obtained from DDA analysis, and a high coincidence rate of 93.2% was observed (Fig. 5e and 5f). More importantly, 1608 of extra proteins (*n* = 8069 vs. *n* = 6461) that were not present in the DDA result were identified. Reducing the information of redundant peptides in the library can downsize the library, thus enabling to increase the efficiency of peptide search. So, we further used Deep4D to predict the charge states of peptides, thus removing those entries of peptides with unreasonable charge states. The accurarcy of charge state prediction of Deep4D that trained on the training set of Pep_CCS reached 94.4% on the test set of Pep_CCS (Supplementary Fig. 7). With charge state prediction, the entries of peptides were reduced by about 60% from 4,770,372 to 2,088,134 (Fig. 5d). However, the identified proteins (*n* = 8075) in DIA analysis of HeLa cell sample with this downsized library (Human_theory_S) didn’t decrease, and a slightly improved coincidence rate (93.4%) to DDA result was observed. The coincidence rate of identified proteins with Human_theory and Human_theory_S libraries was about 95% (Fig. 5f). More importantly, more enriched KEGG pathways could be found based on proteome profiling results yielded from the predicted libraries (Fig. 5g). Although the Peaks Online software has its own evaluation algorithm to ensure the reliability of the protein identification, we also applied a target-decoy strategy to access the false discovery rate. We constructed a decoy library Yeast_decoy by using all peptides generated from reviewed proteins in SwissProt S. cerevisiae database and combined Yeast_decoy with Human_theory and Human_theory_S to perform DIA analysis, respectively. The FDR of two libraries were only 0.98 and 0.96%, which indicates the reliability of the identified results with Human_theory and Human_theory_S libraries. The above results illustrate that 4D DIA proteomics based on theoretical prediction libraries can not only accelerate the proteome profiling, but also greatly improve the depth of protein identification.

### High-coverage DIA phosphoproteomics enhanced by Deep4D-based prediction

Encouraged by the performance of Deep4D in DIA-based proteomics, we expect the predicted 4D libraries could also work well in DIA-based phosphoproteomics. To examine the comparability of the predicted libraries to that of the experimental library, we compared them using the phosphorylated proteome of mouse RAW-264.7 cells. We extracted the whole proteome of RAW-264.7 cells and digested it with trypsin, and then enriched the phosphopeptides by titanium dioxide. The DDA analysis identified 9169 phosphopeptides with MS1, CCS, RT and MS/MS information, which were used to construct the experimental 4D library (RAW_Phos_DDA). Then, four separate phosphopeptides libraries were generated by replacing one or all the information of CCS, RT and MS/MS in RAW_Phos_DDA with the predicted values obtained from Deep4D and DeepPhospho (Supplementary Fig. 8). These five libraries were used for the analysis of DIA data of phosphopeptides from RAW-264.7 cells and compared. As shown in Fig. 6a and 6b, the number of identified peptides and proteins decreased a little bit after replacing the experimental values with the predicted values. When the analysis was performed with the full predicted library, the number of identified phosphorylated peptides and proteins was reduced by only 5.9% and 3.1%, respectively, compared to the experimental library. The results indicate that DIA analysis of phosphopeptides using the prediction library is feasible and that the information of phosphorylated peptides predicted by Deep4D is reliable.

**Fig. 6.**
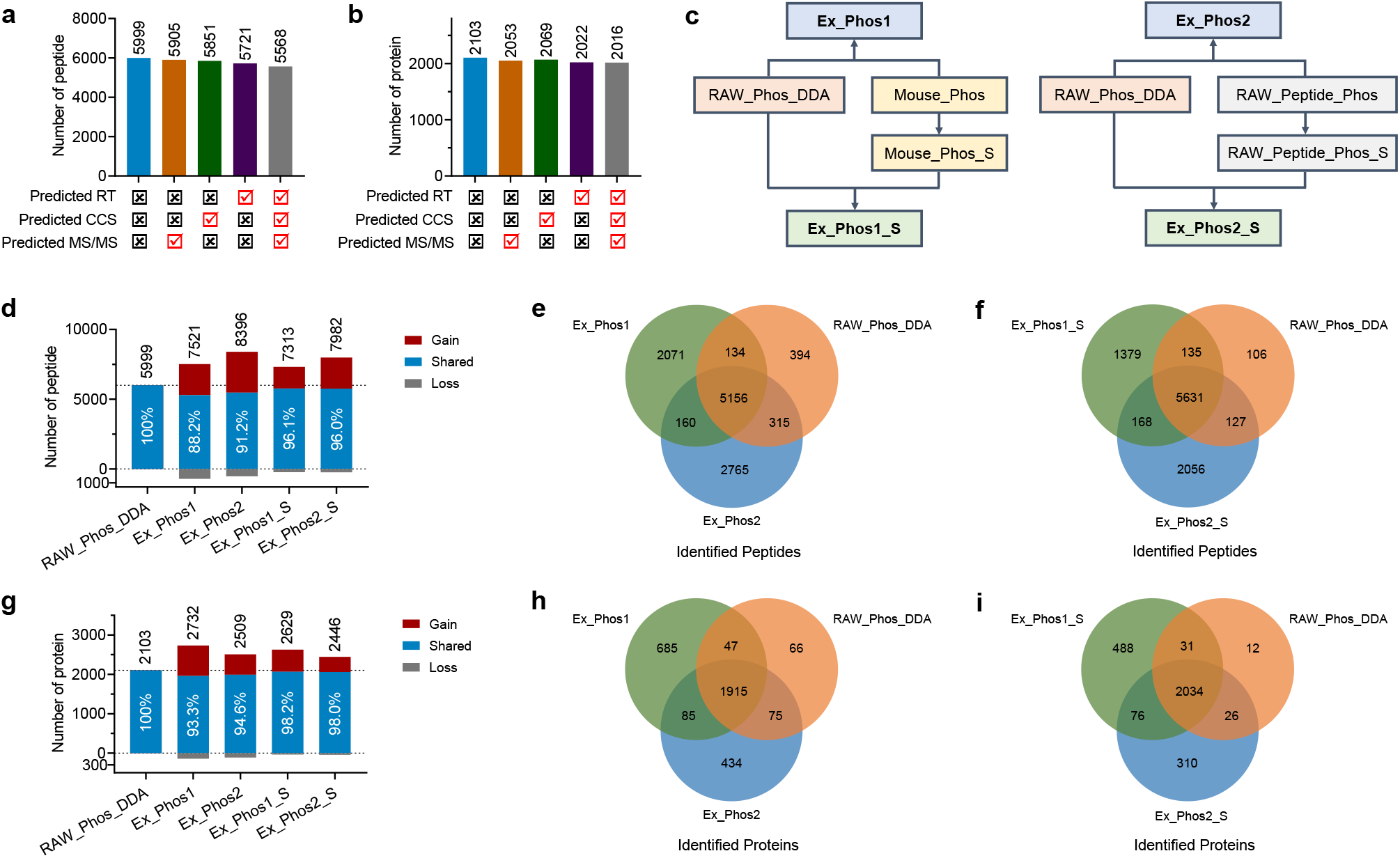
Indeepth DIA-based phosphoproteomics driven by extended 4D libraries. **a-b,** Comparison of the identified numbers of **a**) peptides and **b**) proteins in DIA phosphoproteomics analysis of RAW-264.7 cells with different 4D libraries. **c**, Construction of the extended 4D libraries for DIA phosphoproteomics. **d-f**, Comparison of the identified numbers of phosphopeptides and in DIA analysis of RAW-264.7 cells between extended and experimental 4D libraries. **g-h**, Comparison of the identified numbers of phosphoproteins and in DIA analysis of RAW-264.7 cells between extended and experimental 4D libraries.

We next explore how phosphopeptide prediction libraries can be used to improve the coverage of DIA phosphoproteomics. Different from unmodified proteomics, phosphoproteome owns a more diverse universe which can be reflected in the difference of modification sites and quantity. Due to the excessive number of possible combinations, it is difficult to build a practical predicted library of phosphopeptides from a proteome-wide database at this stage. So, we constructed two extended phosphopeptide 4D libraries (Ex_Phos1 and Ex_Phos2) by combining the experimental phosphopeptide DDA library (RAW_Phos_DDA) with additional phosphopeptide prediction libraries (Fig. 6c, Supplementary Table 4). The Ex_Phos1 is a library consisting of peptides in RAW_Phos_DDA and 540k of theoretical phosphopeptides (Mouse_Phos) derived from a public mouse phosphoproteome database (PhosphoSitePlus), whereas Ex_Phos2 is a library consisting of peptides in RAW_Phos_DDA and 300k simulated phosphopeptides (RAW_Peptide_Phos) derived from the identified unmodified peptides in DDA proteomics results of RAW-264.7 cell sample. As shown in Fig. 6d, more phosphopeptides were identified using the extended phosphopeptide libraries than using only the RAW_Phos_DDA experimental library. 7521 and 8396 phosphopeptides were identified in RAW-264.7 cell sample by using Ex_Phos1 and Ex_Phos2 libraries, respectively, with 88.2% and 91.2% coincidence rate with the results obtained using the experimental library (Fig. 6e).

At the level of protein identification, more proteins were also identified using the extended library than using the experimental library only, with an increase in the number of identifications by 30.0% (Ex_Phos1) and 19.3% (Ex_Phos2), respectively, and an increase in the coincidence rate with the experimental library results to 93.3% and 94.6%, respectively (Fig. 6g and 6h). We also constructed two reversed libraries of Ex_Phos1 and Ex_Phos2 to additionally assess the FDR of peptide identification with the target-decoy strategy. The FDR of Ex_Phos1_decoy and Ex_Phos2_decoy (Supplementary Table 4) were only 0.89% and 0.01%, respectively. These results not only show that the depth of phosphopeptide identification can be greatly improved with the addition of a prediction library, but also demonstrate the high specificity of this strategy. The simulation and prediction of phosphopeptides using unmodified peptides from the same sample can effectively complement the library obtained from DDA phosphopeptide analysis and identify many phosphorylated peptides that have been overlooked.

Given that a downsized library may enhance the analysis efficiency of the DIA data, we adopted an iterative search^17^ approach to reduce the sizes of the two extended libraries. The predicted phosphopeptide libraries Mouse_Phospho or RAW_Peptide_Phos were first applied to the DIA analysis of RAW-264.7 cells to identify phosphopeptides, then new prediction libraries were generated by retaining the identified phosphopeptides (FDR of peptide-to-spectrum match (PSM) less than 3%), and finally these two prediction libraries were combined with the experimental phosphopeptide library (RAW_Phos_DDA) to generate the downsized libraries (Fig. 6c, Ex_Phos1_S and Ex_Phos2_S, Supplementary Table 4). With the two new libraries, the coincidence rate with the experimental library at peptide level was improved to 96.1% and 96.0% (Fig. 6d), and the coincidence rate at protein level was improved to 98.2% and 98.0% (Fig. 6g). It is worth mentioning that only 168 peptides and 76 proteins in the gained part of two libraries coincide, which should be ascribed to the large difference between the two introduced prediction libraries (Fig. 6f and 6i). The result also indicates that it is not adequate to rely only on the sample-specific experimental library for the DIA analysis of phosphopeptides, and many unidentified peptides need to be captured by establishing extended prediction libraries. The strategy we established to combine the prediction library and the experimental library will contribute to the in-depth phosphoproteomics analysis.

## Discussion

4D DIA proteomics is a very promising technique that adds ion mobility dimension to chromatography and mass spectrometry separation, allowing for highly sensitive and reliable analysis and identification. Especially for phosphopeptides, ion mobility separation can distinguish isomers with different site modifications and reduce the complexity of DIA mass spectra.^12^ In this study, we developed a deep learning model based on a self-attention mechanism to establish a 4D DIA proteomics workflow based on in silico prediction library, which overcomes the problems of time consumption and limited identification depth caused by the current DIA analysis mainly relying on experimental library.

Although deep learning-based methods are already available for peptide CCS value and RT prediction, we found that by adding a self-attention module and using a self-attention plus CNN model in the overall framework, one can obtain more accurate results than the state-of-art prediction models. In particular, the prediction of phosphopeptide CCS values has not been achieved yet. We have achieved a highly accurate prediction of phosphopeptide CCS values by constructing a large-scale experimental library, which provides a basis for 4D DIA phosphoproteomics analysis.

A critical limitation of current DIA proteomics methods is that the identification and search range of peptides are limited by the experimental library built based on DDA analysis, which cannot demonstrate the full performance of DIA analysis. By establishing a large-scale 4D theoretical prediction library of peptides from human samples, we have identified 25% more protein than using experimental libraries in the analysis of HeLa cells. In addition, when performing 4D DIA phosphoproteomics analysis, we also found that combining the 4D theoretical prediction library with the DDA experimental library could greatly improve the depth of identification of phosphopeptides and phosphorylated proteins. Taking the analysis of RAW-264.7 cell samples as an example, the amount of identified phosphopeptides and phosphorylated proteins can be increased by 22%-40% and 16%-30%, respectively, using this extended 4D library compared to using the experimental library alone. Therefore, the construction of in silico libraries of peptides by multidimensional prediction can overcome the limitations of experimental libraries and greatly improve the proteome coverage of DIA identification. The established Deep4D model and 4D DIA proteomics and phosphorylation proteomics workflows are expected to be widely applied in proteomics analysis.

## Methods

### Research compliance

All animal experiments and procedures were approved by the committee of experimental animals of Wuhan University. All experimental protocols and animal handling procedures were performed in accordance with the National Institute of Health Guide for the Care and Use of Laboratory Animals (NIH Publications No. 80-23) revised 1996 and the experimental protocols were approved by the committee of experimental animals of Wuhan University.

### Sample preparation

The HeLa cells, HepG2 cells, RAW-264.7 cells and Yeast were all cultivated following standard protocols. For the preparation of peptide samples, the urea lysis buffer solution (8 M urea) was first added to the cells or tissue homogenates, and the mixture solutions were subsequently sonicated in cold water bath (Cheng Xian, China). Then, the sample lysates were centrifuged at 12,000 *g* for 20 min at 4 °C. The supernatant was collected for subsequent protein quantification and digestion. Extracts from each sample was reduced with 10 mM dithiothreitol at 37 °C for 45 min and alkylated with 10 mM iodoacetamide at room temperature in the dark for additional 30 min. Then ammonium bicarbonate was added to dilute the urea concentration to 1 M. Proteolysis was performed overnight at 37 °C by adding trypsin in a 1:50 enzyme:protein (wt/wt) ratio and then desalted using Sep-Pak tC18 Cartridges (Waters, USA). TiO_2_ (Thermo Fisher Scientific, USA) was applied to enrich phosphopeptides. Due to the limited quantities, peptides of E.coli and Drosophila melanogaster were not used for phosphopeptides enrichment. All peptide solutions were dried by Centrifugal Vacuum Concentrators (labconco, USA) and reconstituted in 0.1% formic acid for further analysis.

### LC-MS/MS analysis

All the LC-MS analysis of peptides were performed on timsTOF Pro mass spectrometer (Bruker Daltonics, Germany) with DDA or DIA modes *via* a CaptiveSpray nano-electrospray ion source (Bruker Daltonics, Germany) coupled with an EASY-nLC 1200 system (Thermo Fisher Scientific, USA). Peptides were loaded on a 25 cm in-house packed HPLC-column (75 μm inner diameter packed with 3 μm Venusil XBP C18 silica beads, Agela Technologies, China). The mobile phase for gradient elution consisted of mobile phase A (ddH2O mixed with 0.1% formic acid) and mobile phase B [ACN-H2O (8:2, v/v) mixed with 0.1% formic acid]. The optimal chromatographic gradient program for both DDA and DIA analysis was set as follows: 2~28% B at 0~90 min, 28~46% B at 91~100 min, 46~95% B at 101~110 min, 95% B at 111~120 min. A shortened gradient of 60 min (2~28% B at 0~45 min, 28~46% B at 46~50 min, 46~95% B at 51~55 min, 95% B at 56~60 min) was also used for some samples. The m/z of ion and the ion mobility were calibrated linearly using three ions from the Agilent ESI LC/MS tuning mix (m/z, 1/K_0_: 622.0289, 0.9848 Vs·cm^-2^; 922.0097, 1.1895 Vs·cm^-2^; and 1221.9906, 1.3820 Vs·cm^-2^).

For DDA analysis, the timsTOF Pro mass spectrometer was operated in data-dependent PASEF mode with 1 survey TIMS-MS and 10 PASEF MS/MS scans per acquisition cycle. Ion accumulation and ramp time in the dual TIMS analyzer was set to 100 ms each, and the ion mobility range was from 1/K_0_ = 1.6 Vs·cm^-2^ to 0.6 Vs·cm^-2^. The collision energy was lowered linearly as a function of increasing mobility starting from 59 eV at 1/K_0_ = 1.6 Vs·cm^-2^ to 20 eV at 1/K_0_ = 0.6 Vs·cm^-2^. Precursor ions for MS/MS analysis were isolated with a 2 Th window for m/z < 700 and 3 Th for m/z >800 in a total m/z range of 100-1700 by synchronizing quadrupole switching events with the precursor elution profile from the TIMS device. Singly charged precursor ions were excluded with a polygon filter (otof control, Bruker Daltonik GmbH). Precursors for MS/MS were picked at an intensity threshold of 2500 arbitrary units (a.u.) and re-sequenced until reaching a ‘target value’ of 20000 a.u. considering a dynamic exclusion of 24 s elution.

For DIA analysis, the mass spectrometer was operated in positive ion data-independent acquisition (diaPASEF) mode. The capillary voltage was set to 1400 V. The MS and MS/MS spectra were acquired from 100 to 1,700 m/z with an ion mobility range (1/K_0_) of 0.6 Vs·cm^-2^ to 1.6 Vs·cm^-2^. The ramp and accumulation time were set to 100 ms to achieve a duty cycle close to 100%. Polygonal filtering was applied to exclude low m/z and singly charged ions for precursor selection. The collision energy was ramped linearly as a function of mobility from 59 eV at 1/K_0_ = 1.6 Vs· cm^-2^ to 20 eV at 1/K_0_ = 0.6 Vs·cm^-2^. To perform diaPASEF acquisition mode, we defined 25 Th isolation windows from m/z 400 to 1,200 and totally 64 windows in the DIA experiment.

### Construction of large-scale phosphopeptide CCS database

The phosphopeptides prepared from HeLa cells, HepG2 cells, tissues of brain, kidney and liver of mouse, and Yeast were analyzed at DDA mode with 120 min LC gradient each with three technical replicates. The peptides identification was performed by Maxquant^23^ (version 1.6.15.0), which could extract 4D information and MS/MS spectra of ions from MS raw files. The phosphorylation of serine, threonine and tyrosine, oxidation of methionine and acetylation on N-terminal of proteins were chosen as variable modifications and carbamidomethylation of cysteine was set as fixed modifications. Semi-specific and specific digestion modes were performed, respectively. The maximum number of modifications per peptide was set to 5, while the maximum peptide mass and minimum peptide length were set to 4600 Da and 7, respectively. For semi-specific mode, minimum and maximum length were set to 8 and 25. The false discover rate (FDR) of peptide-spectrum match (PSM) was set to 0.01. Maximum precursor and fragment ion mass tolerance were searched as default for TIMS-DDA data. The FASTA files of reviewed reference proteomes for each organism were downloaded from UniProt (https://www.uniprot.org).

After finishing the peptide identification, the doubly and triply phosphopeptides were selected. The CCS of peptides in each sample was calibrated by the calibration curve which was calculated in DataAnalysis (version 5.3, Bruker) according to the ion mobility of three ions from the Agilent ESI LC/MS tuning mix. Even a peptide with the same charge state may adopt multiple conformations in gas phase, so we keep the CCS of one peptide that has the highest intensity. To acquire high confidence phosphopeptide, the modification probability threshold was set at 0.95 for phosphosite. Then all peptides were integrated and RSD of each peptide’s CCS were calculated and the peptides which had RSD over 0.4% would be deleted. Finally, we combined semi-specific with specific results to constructed a high confidence dataset containing CCS values of 44,424 doubly charged and 9,701 triply charged phosphopeptides.

### Deep4D model

#### Peptide representation

We firstly convert peptide into a letter sequence. 20 kinds of amino acid are represented according to the capital letter, like Alanine as “A”, Lysine as “K”, and “a” will be put in the beginning of the sequence if there is acetylation on N-terminal of peptide. Then we use “s”, “t”, “y” to represent the phosphorylation of serine, threonine, tyrosine, and “e” represent oxidation of methionine. Each amino acid was coded into a 23 dimensional one-hot vector. The first twenty dimensions of the vector represent 20 kinds of amino acids, and the last three dimensions represent acetylation, oxidation and phosphorylation. The maximum length of peptide was set to 49. For a peptide containing M amino acids, the M × 23 one-hot tensors will be padded with zero to 50 × 23. To make use of the position information of the sequence, we injected some information about the absolute position of one-hot tensor through a “positional encodings”.^14^ Then the one-hot tensor and “position encoding” were added to form the representation tensor of peptide with 50 × 23 dimention.

#### Model architecture

The deep learning model was written by Python with Pytorch. Considering the sequential characteristic of peptides, the encoder-decoder structure was chosen, while it was not the usual “sequence to sequence” but “sequence to one”.

##### (1) Encoder

The representation tensor was firstly embedded through a 500-dimensional full connected layer. Then the embedding tensor was put into several serial sub-encoders, two sub-encoders for ion mobility prediction and three sub-encoders for retention time prediction. Each sub-encoder consists of 5 layers, and each layer was composed of multi-head attention and fully connected feed-forward network, with residual connection and layer normalization. The number of heads and dimension of feed-forward network was set to 5 and 1200.

##### (2) Decoder

The decoder consists of convolutional neural network and fully connected feed-forward network. The output tensor of Encoder was put into convolutional neural network. For ion mobility prediction, the convolutional neural network contains 2 batch normalization layers, 2 convolution layers and 2 max pooling layers. For retention time prediction, the convolutional neural network contains 3 batch normalization layers, 3 convolution layers and 3 max pooling layers. Then, the 3D output tensor of convolutional neural network was converted to 1D and put into a 4 layers fully connected feedforward network to obtain the final predicted ion mobility or retention time.

#### Model training

We trained or fine-tuned our models on a machine with one NVIDIA Tesla V100 GPU. The Adam optimizer was used with beta1= 0.9, beta2 = 0.999 and eta_min = 10-9. Learning rate was varied by the CosineAnnealingWarmRestarts method.

For CCS prediction of unmodified peptides, we used three models to predict doubly, thriply and quadruply peptide, respectively. We pretrain Deep4D based on a dataset: DeepCCS, which consists of many unmodified peptide CCS acquired from several different species. Deep4D was firstly trained on the training set contains 333,079 unique CCS value of doubly charged peptides. For the triply charged and quadruply charged peptides, the model of doubly charged peptides were fine-tuned with 112,905 triply charged peptides CCS and 19,572 quadruply peptides CCS, respectively.

For CCS prediction of phosphopeptides, we randomly divided the phosphopeptide dataset into train and testing sets. The number of train and testing set of doubly charged phosphopeptides are 39,978 and 4,446. The number of train and testing set of triply charged phosphopeptides are 8,730 and 971. We then fine-tuned the pre-trained Deep4D with douly and triply charged train sets, respectively.

For RT prediction, we used one model to predict all peptides. Like the training of ion mobility model, we also pretrained Deep4D on a dataset: SWATH, which contains retention time of more than 140k normal peptides. Then we fine-tuned Deep4D with other datasets.

#### Comparing Deep4D with other models

To avoid possible unfairness in retraining others model, here, we only compared Deep4D with others models on the training and testing sets mentioned in their works. We also used the same metrics to compare our model’s performance with their models’ performance reported in their article. For CCS prediction, we compared Deep4D with the LSTM model proposed by Meier et al. We trained Deep4D with the training set mentioned above and tested Deep4D on the testing set containing 18,5248 synthetic peptides from ProteomeTools library.^13^ Median relative error was used as metric to measure the performance of model. For retention time prediction, we compared Deep4D with DeepLC on 15 unmodified peptide datasets. We firstly train and test Deep4D on a dataset (SWATH library), and then fine-tuned model with other 14 datasets. Prediction performance was measured by mean absolute error and Δt95%. Then we compared Deep4D with DeepPhospho on three phosphopeptide retention time datasets: RPE1 DDA, RPE1 DIA and U2OS DIA. We firstly train and test Deep4D with RPE1 DDA, and then fine-tuned with RPE1 DIA and U2OS DIA. Median absolute error was used to measure the performance.

#### Charge state prediction

Here we used a three dimensional one-hot vector to represent charge +2, +3, and +4. The dimension last fully connected layer was change from 1 to 3 and Sigmoid function was added at the end of the model. We pre-trained the model based on the dataset Pep_CCS and fine-tuned it with current DDA result to get a instrument-specific model.

### Generation of 4D libraries DIA-based proteomics

#### DIA proteomics

Firstly, Deep4D and DeepPhospho were fine-tuned with the DDA results of RAW-264.7 proteome. Four libraries with the same peptide list as HeLa_DDA were generated with fine-tuned Deep4D and DeepPhospho. The peptide list of Human_theory was generated by using Protein Digestion Simulator (version 2.4.7993) to carry out tryptic specific digestion of human proteome from SwissProt H. sapiens database with missed cleavage set to 2 and mass range set from 500 to 6800. These peptides were finally filtered by set length range from 7 to 49 and given charge 2 and 3 to form the peptide list of Human_theory. The downsizing peptide list of Human_theory_S was generated with charge state prediction model. Finally, each library was built based on its peptide list by fine-tuned Deep4D and DeepPhospho.

#### DIA phosphoproteomics

For the four libraries with the same peptide list as RAW_Phos_DDA, Deep4D were fine-tuned with the unmodified peptides in DDA results of RAW-264.7 phosphoproteome, and DeepPhospho were fine-tuned with the DDA results of HeLa cell phosphoproteome. For Ex_Phos1 and Ex_Phos2, Deep4D and DeepPhospho were fine-tuned with the DDA results of RAW-264.7 phosphoproteome. The peptide list of RAW_Phos_DDA was got from phosphopeptides with Ascore > 10 in the DDA results of RAW-264.7 phosphoproteome. The peptide list of Ex_Phos1 was generated by combining RAW_Phos_DDA with tryptic specific digestion of Mouse phosphoproteome from PhosphoSitePlus with missed cleavage set to 2 and length range set from 7 to 49. The peptide list of Ex_Phos2 was generated by combining RAW_Phos_DDA with all the possible mono-phosphopeptide based on the unmodified peptide in DDA results of RAW-264.7 proteome. Each phosphopeptide in Ex_Phos1 and Ex_Phos2 was given charge 2 and 3. All the libraries were built based on their peptide list with fine-tuned Deep4D and DeepPhospho. Ex_Phos1_S and Ex_Phos2_S were generated from the identified peptides with FDR of PSM < 3%.

### DIA data analysis and false discovery rate evaluation

PEAKS Online was used to perform DIA data analysis. We set precursor mass error tolerance to 20 ppm and fragment mass error tolerance to 0.05 Da. The CCS error tolerance was set to 0.05. The PSM FDR and −10LgP of proteins was set to 1% and 20, respectively. Reference proteome were all the reviewed proteome of sample’s organism downloaded from Uniprot.

Although the FDR of peptide and protein has been evaluated in the DIA analysis of PEAKS Online, we apply another target-decoy strategy to assess the FDR of predicted 4D library. For DIA analysis of HeLa proteome, we constructed a decoy library Yeast_decoy containing doubly and triply charged yeast tryptic peptides generated from all reviewed proteins in SwissProt S. cerevisiae database. The RT, CCS and fragment intensity of decoy library was predicted by Deep4D and DeepPhospho with the same parameters as the target library. Then we combined Yeast_theory with Human_theory and Human_theory_S to build two target libraries. Each target library was used to do DIA analysis with FDR of PEAKS Online set to 1% on the protein database which combine human and yeast. The FDR of each target library was calculated by D/((D+T)). D represents the number of identified yeast proteins and T represents the number of identified human proteins

For DIA analysis of RAW-264.7 phosphoproteome, the decoy libraries of Ex_Phos1 and Ex_Phos2 were generate by reversing the sequences of all peptides in the target library except the C-terminal residue, and each phosphopeptide was given 2+ and 3+ charge state. The RT, CCS and fragment intensity of decoy library was then predicted by Deep4D and DeepPhospho with the same parameters as the predicted target library. And we searched the DIA data on PEAKS Online based on target library and decoy library, respectively. The FDR was calculated as FDR = D/T, D represents the number of peptide or proteins identified in decoy library, T represents the number of peptides or proteins identified in target library.

## Acknowledgements

This work was financially supported by the National Natural Science Foundation of China [22074111 (S.M.C.)] and National Key Research Development Program of China [2021YFC2700700]. We also thank the support of the start-up funds of Wuhan University (S.M.C.) and the National Youth Talents Plan of China (S.M.C.). The authors thank Guibao Shen for fruitful discussions.

