## Supplementary Information for "High-Coverage Four-Dimensional Data-Independent Acquisition Proteomics and Phosphoproteomics Enabled by Deep Learning-Driven Multi-Dimensional Prediction"

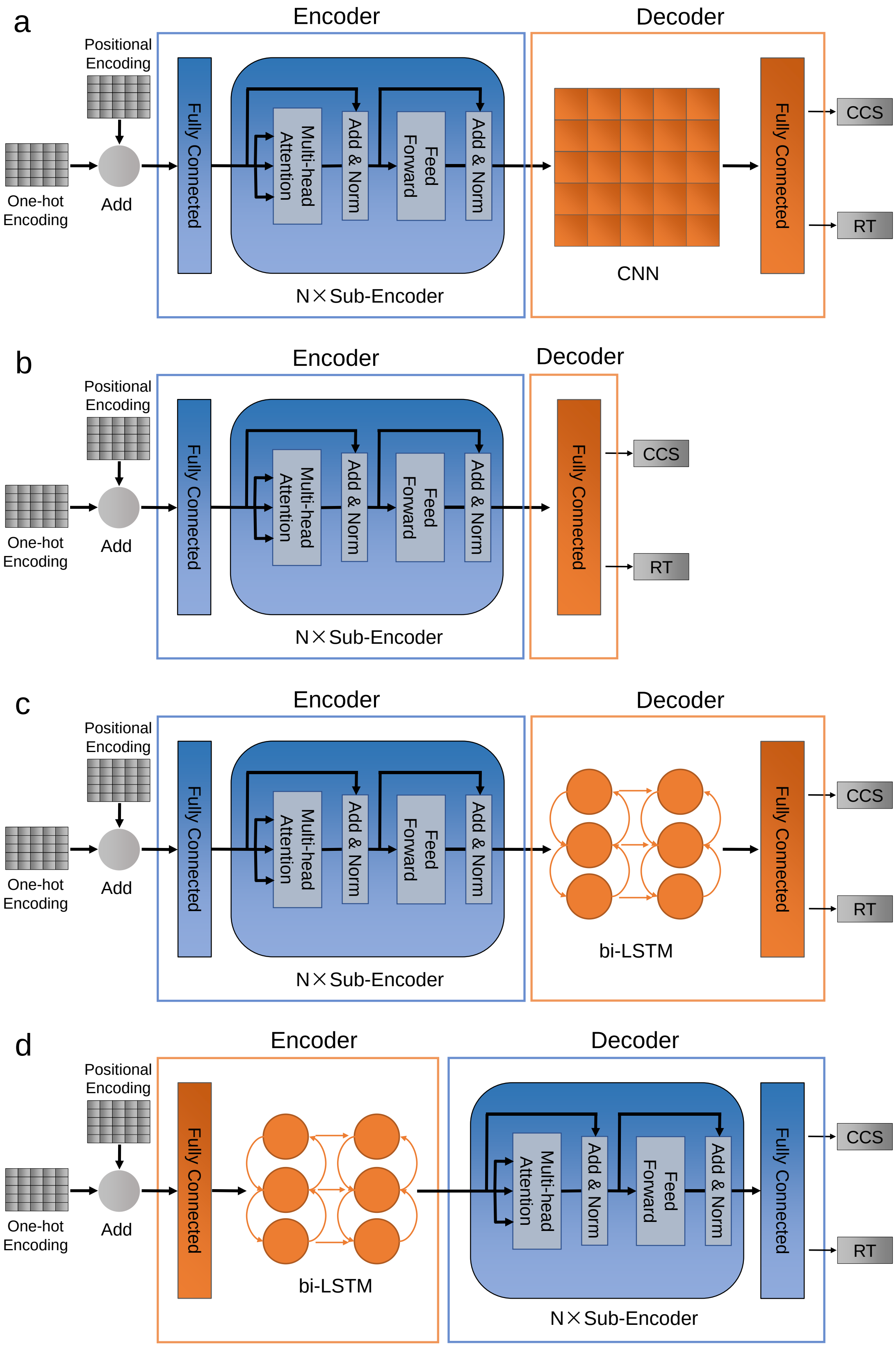


**Supplementary Fig. 1 | The structures of different deep learning models. a** Deep4D (Self-attention + CNN). **b** Self-attention. **c** Self-attention + bi-LSTM. **d** bi-LSTM + Self-attention.

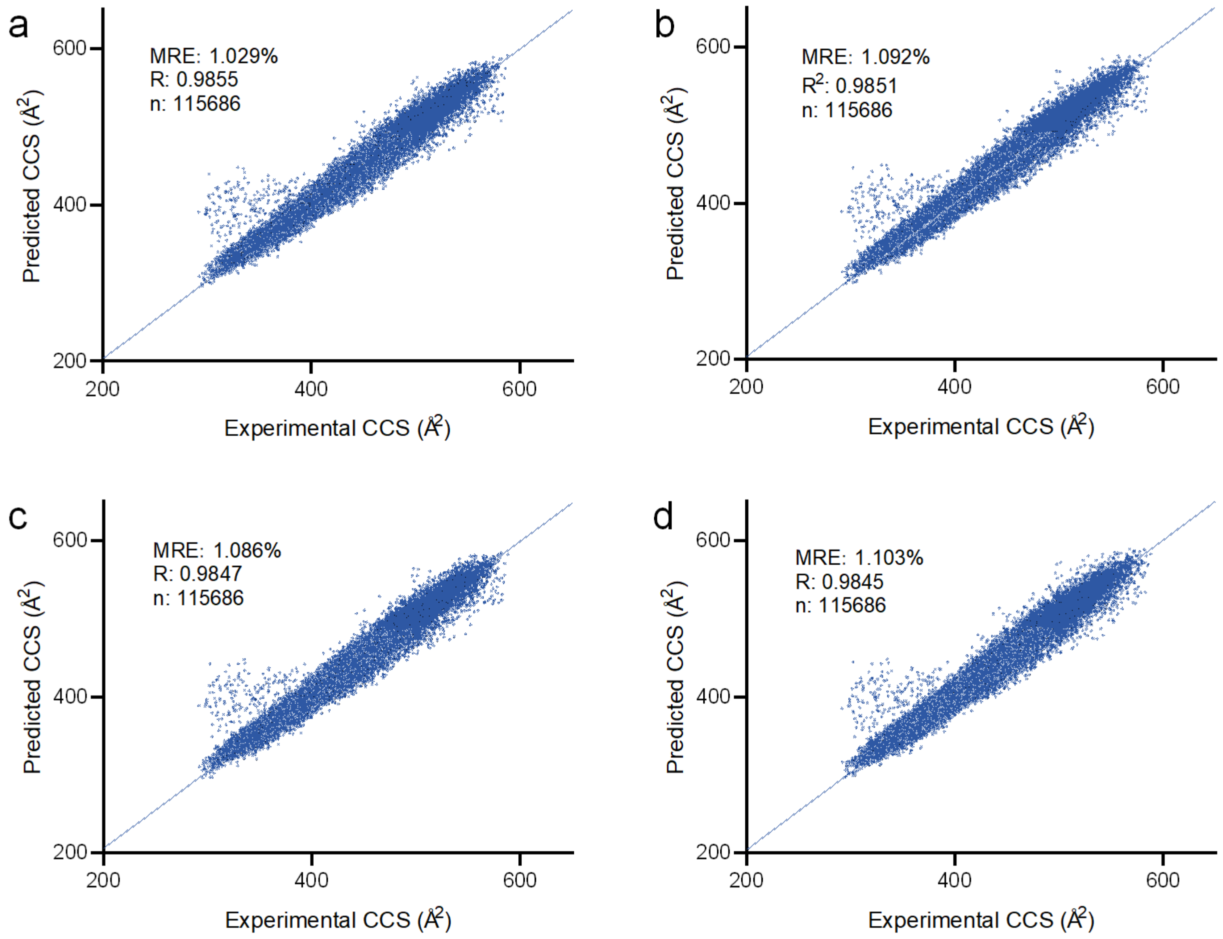


**Supplementary Fig. 2 | Correlation of predicted versus experimental CCS values on the doubly charged peptides derived from PepCCS dataset using different deep learning models. a** Deep4D (Self-attention + CNN). **b** Self-attention + LSTM. **c** Self-attention. **d** LSTM + Self-attention.

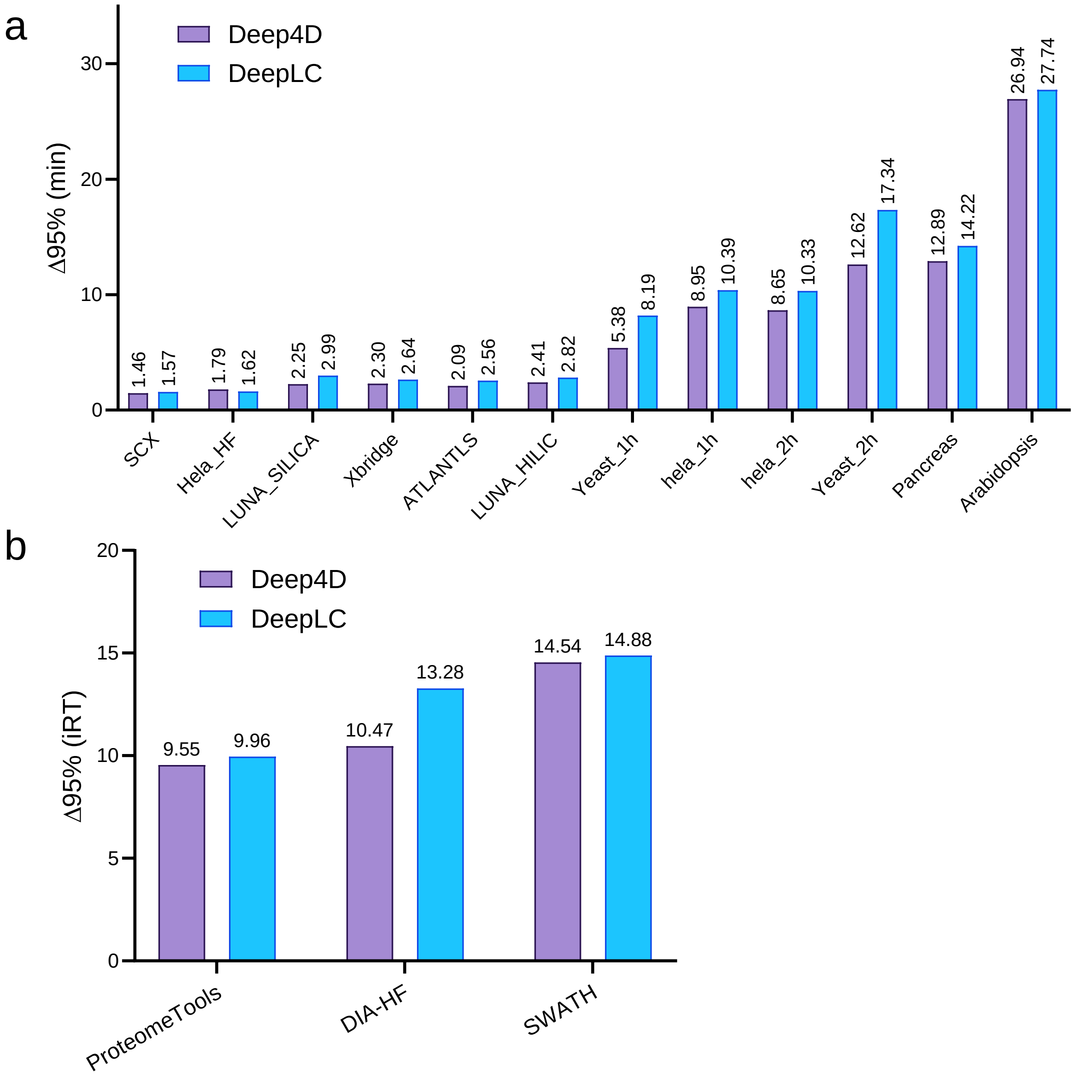


**Supplementary Fig. 3 | ** Comparison of $\Delta t_{95\%}$ of (**a**) RT and (**b**) iRT between Deep4D and DeepLC in RT prediction of 15 datasets, respectively.

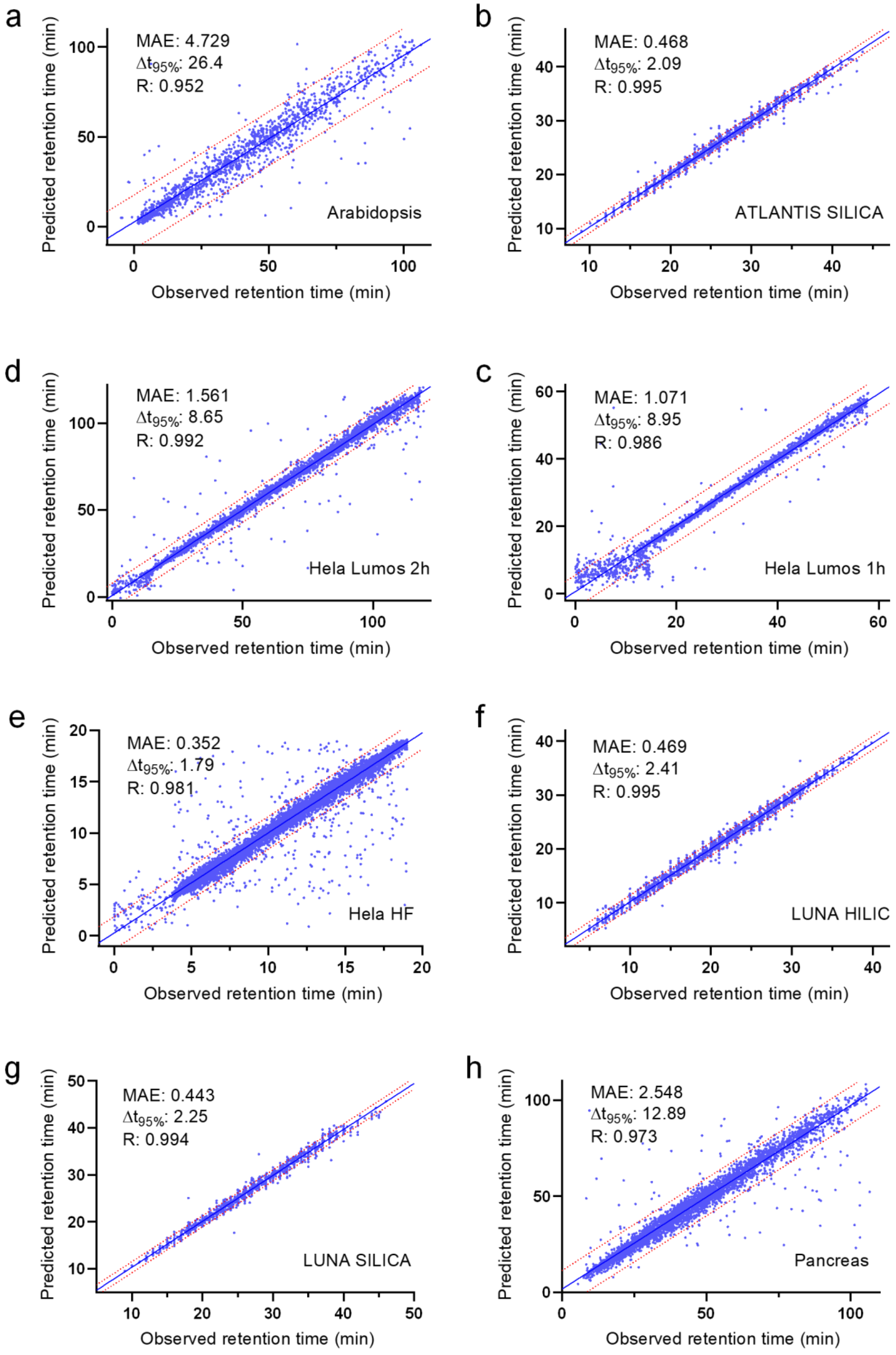
a
MAE: 4.729
Δt95%: 26.4
R: 0.952
Arabidopsis
b
MAE: 0.468
Δt95%: 2.09
R: 0.995
ATLANTIS SILICA
d
MAE: 1.561
Δt95%: 8.65
R: 0.992
Hela Lumos 2h
c
MAE: 1.071
Δt95%: 8.95
R: 0.986
Hela Lumos 1h
e
MAE: 0.352
Δt95%: 1.79
R: 0.981
Hela HF
f
MAE: 0.469
Δt95%: 2.41
R: 0.995
LUNA HILIC
g
MAE: 0.443
Δt95%: 2.25
R: 0.994
LUNA SILICA
h
MAE: 2.548
Δt95%: 12.89
R: 0.973
Pancreas
Predicted retention time (min)
Observed retention time (min)

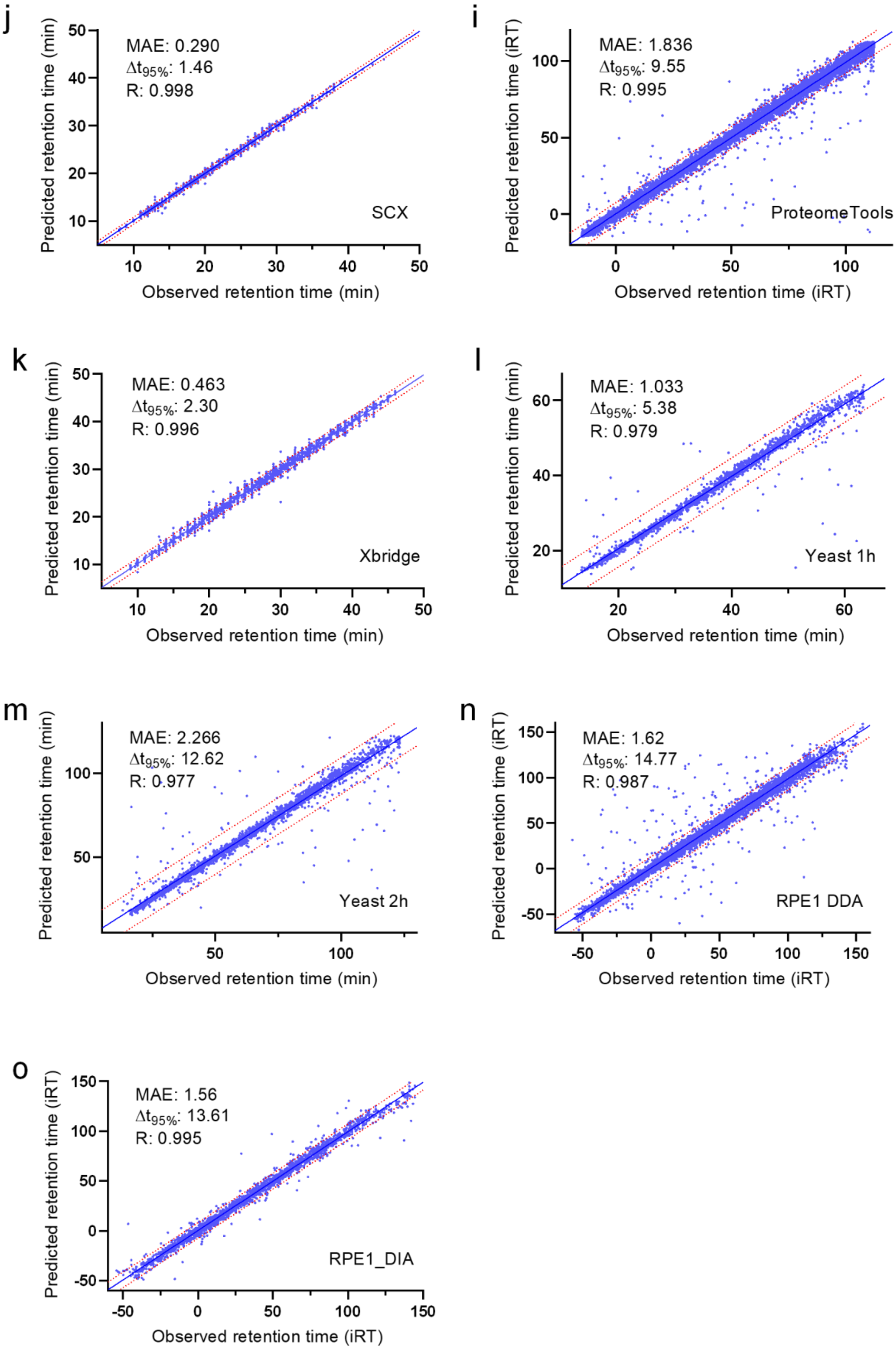


**Supplementary Fig. 4 | a-o** Scatter plots for predicted against observed RT and iRT on the test data respectively. Median absolute error (MAE), $\Delta t_{95\%}$ and Pearson correlation coefficient (R) are indicated.

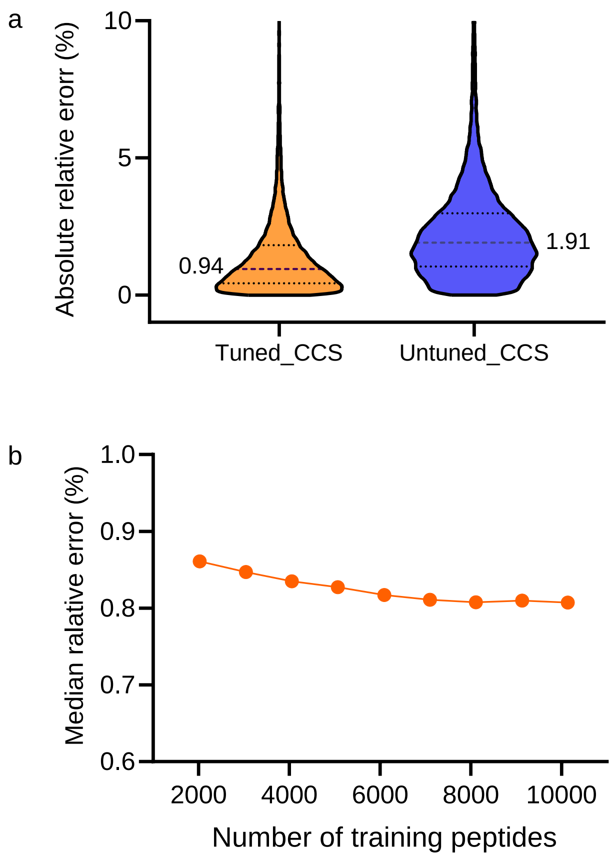


**Supplementary Fig. 5 | a** Distribution of absolute relative error of peptide CCS in Brain1 dataset predicted by fine-tuned and untuned Deep4D model. **b** Relationship between the number of training peptides for Deep4D and its prediction accuracy.

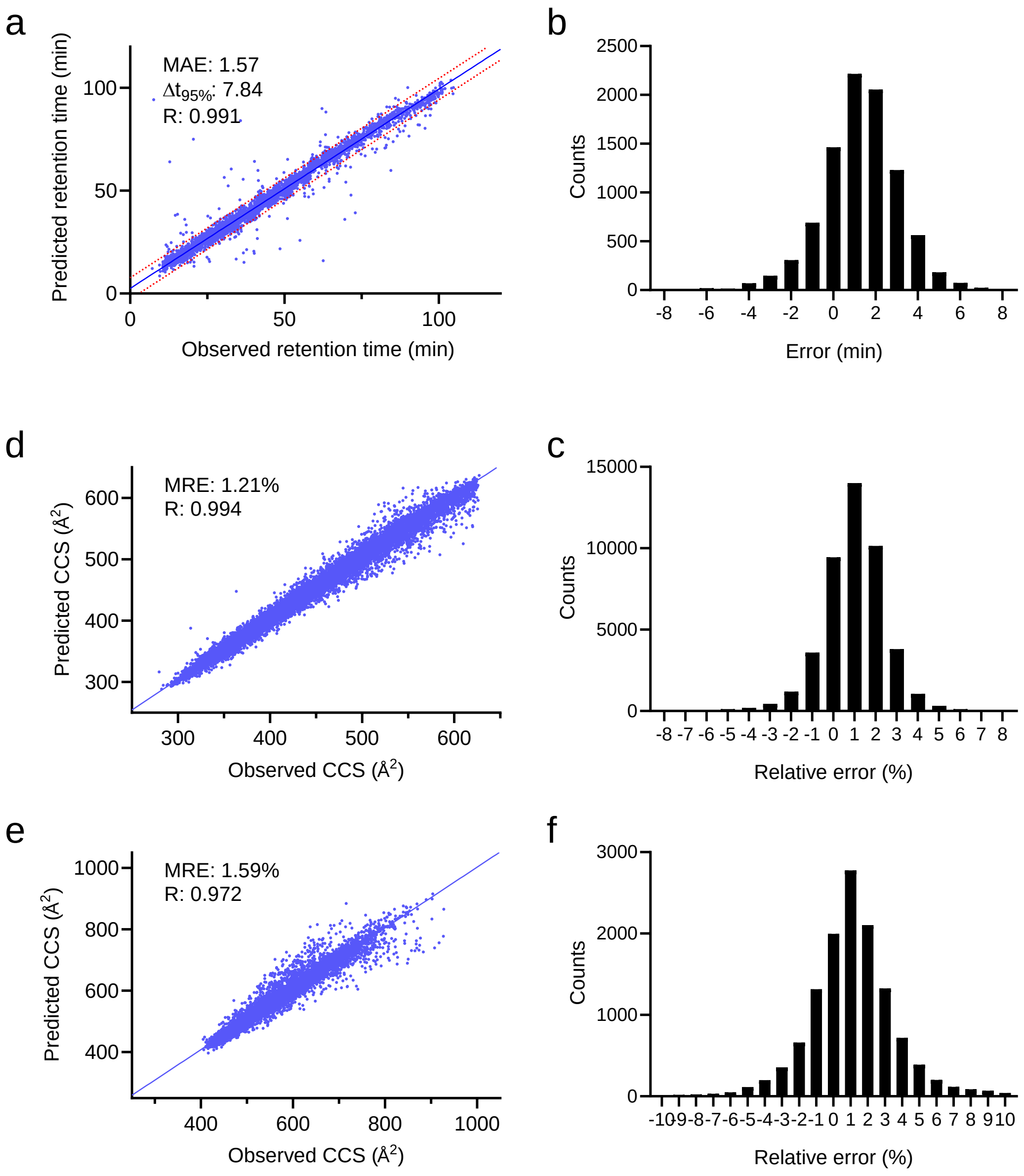


**Supplementary Fig. 6 | a** Correlation of predicted versus experimental RT of peptides with Deep4D on experimental Hela library. **b** Distribution of the error of predicted RT. **c** Correlation of predicted versus experimental CCS values of doubly charged peptides with Deep4D on experimental Hela library. **b** Distribution of the relative error of predicted CCS values of doubly charged peptides. **e** Correlation of predicted versus experimental CCS values of triply charged peptides with Deep4D on experimental Hela library. **f** Distribution of the relative error of predicted CCS values of triply charged peptides.

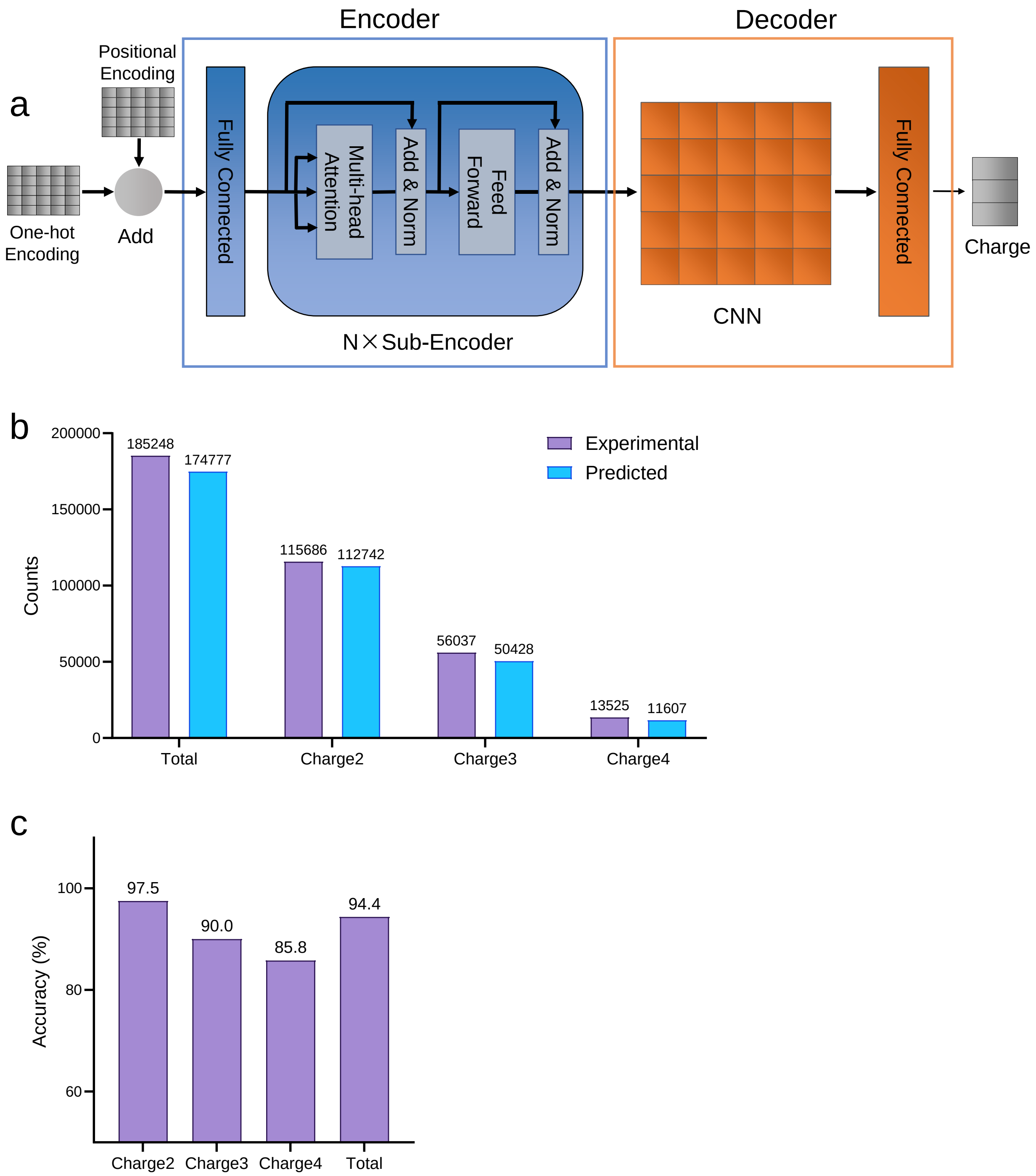


**Supplementary Fig. 7 | a** Deep4D for predicting possible charge states of peptide. The output is a 3-dimention one-hot vector which represents 2+, 3+ and 4+ charge states. **b** The predicted charge state vs. experimental charge state of peptides in the test set of Pep_CCS. **c** The prediction accuracy of each charge state and total peptides.

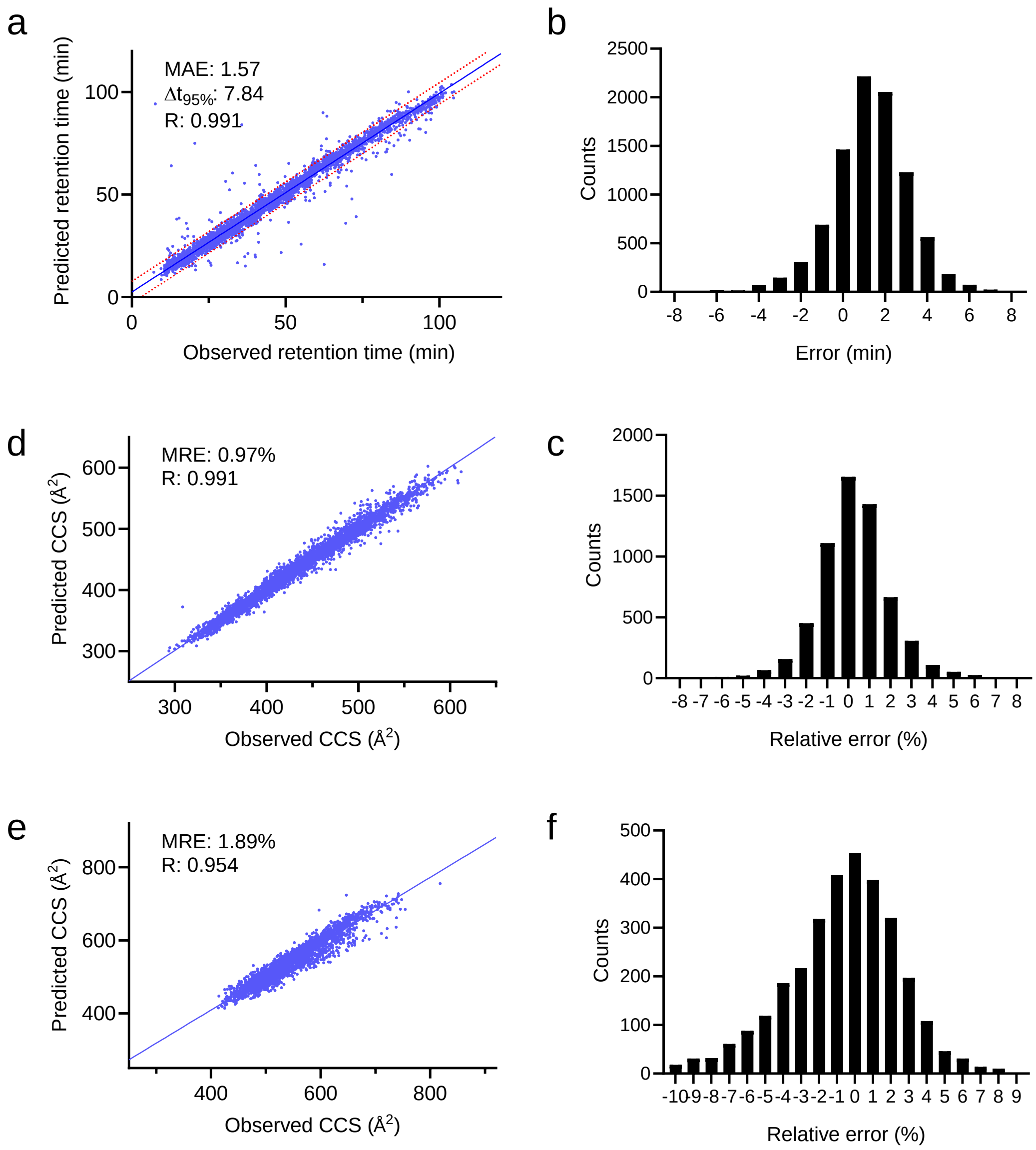


**Supplementary Fig. 8 | a** Correlation of predicted versus experimental RT of peptides with Deep4D on RAW_Phos_DDA library. **b** Distribution of the error of predicted RT. **c** Correlation of predicted versus experimental CCS values of doubly charged peptides with Deep4D on RAW_Phos_DDA library. **b** Distribution of the relative error of predicted CCS values of doubly charged peptides. **e** Correlation of predicted versus experimental CCS values of triply charged peptides with Deep4D on RAW_Phos_DDA library. **f** Distribution of the relative error of predicted CCS values of triply charged peptides.

**Supplementary Table 1 |** Overview of the phosphopeptide CCS dataset by samples and different proteolytic enzymes. The numbers of phosphopeptide in different samples by using trypsin and chymotrypsin proteolysis were shown in the table.

| Samples | Peptide numbers | |
|---|---|---|
| | Trypsin | Chymotrypsin |
| HeLa | 3,290 | 3,904 |
| HepG2 | 10,341 | 7,830 |
| Brain | 11,938 | 5,701 |
| Liver | 6,351 | 4,537 |
| kidney | 9,134 | 5,732 |
| Yeast | NA | 6,425 |

**Supplementary Table 2 |** Overview of the phosphopeptide CCS dataset by number of phosphosites and charge state.

| Phosphate group number | Peptide numbers | |
|---|---|---|
| | Charge State 2+ | Charge state 3+ |
| 1 | 29,654 | 5,240 |
| 2 | 11,889 | 3,236 |
| ≥3 | 2,881 | 1,225 |

**Supplementary Table 3 | ** Overview of the phosphopeptide CCS dataset by amino acids of C-term.

| C-term amino acids | Peptide numbers |
|---|---|
| R | 13358 |
| K | 13344 |
| L | 7470 |
| F | 4908 |
| Y | 3926 |
| M | 1650 |
| D | 1376 |
| W | 1060 |
| N | 1049 |
| Q | 939 |
| S | 899 |
| E | 785 |
| A | 734 |
| H | 617 |
| T | 600 |
| V | 399 |
| I | 280 |
| G | 252 |
| P | 246 |
| C | 233 |

**Supplementary Table 4 | ** The peptide information of different libraries for 4D DIA phosphoproteomic.

| Datasets | Peptide numbers | | |
|---|---|---|---|
| | Peptide entries (2+) | Peptide entries (3+) | Total |
| RAW_Phos_DDA | 6,085 | 3,084 | 9,169 |
| Ex_Phos1 | 278,322 | 275,321 | 553,643 |
| Ex_Phos2 | 159,831 | 156,830 | 316,661 |
| Ex_Phos1_S | 13,870 | 10,869 | 24,739 |
| Ex_Phos2_S | 13,133 | 10,132 | 23,265 |
| Ex_Phos1_decoy | 272,237 | 272,237 | 544,474 |
| Ex_Phos2_decoy | 153,746 | 153,746 | 307,492 |
